# Disrupted salience network dynamics during the imagery of migraine attacks

**DOI:** 10.1101/2025.05.10.653239

**Authors:** Inês Esteves, Alexandre Perdigão, Ana R. Fouto, Amparo Ruiz-Tagle, Gina Caetano, Joana Cabral, Isabel Pavão Martins, Raquel Gil-Gouveia, César Caballero-Gaudes, Patrícia Figueiredo

## Abstract

Moderate to severe head pain is a hallmark of recurring migraine attacks. However, it is challenging to study patients during spontaneous attacks and most research on the brain mechanisms of pain in migraine patients has been limited to the processing of painful stimuli between attacks. Here, we hypothesize that the experience of a migraine attack extends beyond the response to painful stimuli and is associated with specific impairments of the salience network (SN), which integrates sensory, emotional and cognitive information in relation to salient stimuli. To test this hypothesis, we analysed the SN dynamics of a group of patients with episodic migraine in three distinct conditions: at rest during a spontaneous migraine attack (ictal phase); while performing an imagery task aiming to elicit the experience of a previous attack, during the interictal phase; and at rest, during the interictal phase. For comparison, we also studied a group of healthy controls in three matching conditions, including rest as well as an imagery task of a (non-migraine) head pain experience. We collected functional magnetic resonance imaging (fMRI) data and used a dynamic functional connectivity (dFC) analysis to examine the temporal features of the SN from a total of 78 samples. Compared to healthy controls, the SN had a significantly shorter lifetime in patients during the pain imagery task, but not during a migraine attack or interictal resting state. Our results support the disruption of the SN in migraine, and indicate that pain imagery may be a useful paradigm for isolating the emotional and cognitive aspects of pain and investigating SN dynamics.

## 1. Introduction

Migraine is a cyclic neurovascular disorder marked by recurrent episodes of pulsating headaches and an increased sensitivity to sensory stimuli, leading to symptoms such as photophobia, phonophobia, and nausea (Goadsby et al., 2017). The active headache period, known as the ictal phase, typically lasts between 4 and 72 hours (Peng & May, 2020), while the interictal phase refers to the intervals between attacks. Functional magnetic resonance imaging (fMRI) studies measuring responses to visual, olfactory, and painful stimuli have consistently shown that migraine patients exhibit abnormal sensory processing, lack typical habituation between attacks, and display atypical connectivity in sensory networks (de Tommaso et al., 2014; Russo et al., 2018; Schwedt et al., 2015). This persistent sensitivity to sensory and painful stimuli is closely linked to the function of the salience network (SN), which is crucial for detecting, filtering, and prioritizing relevant information (Seeley, 2019). Anchored in the anterior insula and anterior cingulate cortex (ACC), it acts as a switch between self-focused and externally-focused brain networks, regulating attention, and integrating sensory, emotional, and cognitive information (Schimmelpfennig et al., 2023). Thus, disruptions within this network may contribute to the increased sensitivity to sensory stimuli and the deficits in sensory processing seen in individuals with migraine both during the attacks and between them. Consistent with this idea, neuroimaging studies have frequently reported alterations in the SN of migraine patients (Xue et al., 2012; D. Yu et al., 2017), as well as particularly within both the ACC (Z. Li et al., 2016; Liu et al., 2023; Ou et al., 2024; Wang et al., 2022; D. Yu et al., 2017) and the insula (Araújo et al., 2023; Coppola et al., 2018; Huang et al., 2024; Lim et al., 2021; Tian et al., 2021; Wei et al., 2020; Z. Yu et al., 2017).

Changes of the SN may be indicative of impaired cortical excitability (Coppola et al., 2019). These changes could reflect disproportionate responses in hyperexcitable regions when exposed to external stimuli, a pattern that aligns with migraine’s characteristic difficulties in adapting to sensory input and its disrupted balance of excitatory and inhibitory neurotransmitters (Aurora & Wilkinson, 2007; O’Hare et al., 2023; Vecchia & Pietrobon, 2012; Younis et al., 2017). To assess this, <u>Veréb et al. (2020</u>) studied the dynamic functional connectivity (dFC) of the SN of migraine patients, both with and without aura, during the interictal phase. Although the study only observed changes in migraine patients with aura, increased cortical excitability is also a feature of migraine without aura (Cortese et al., 2017). Consequently, we propose that this phenomenon is present and may be elicited during pain imagery experiences in migraine patients without aura.

Previous studies have mainly examined the SN’s functional connectivity in the interictal phase, either during rest or in response to painful stimulation. Here, we hypothesize that involvement of the SN in the experience of a migraine attack is distinct from other types of pain experience. To address this, we aim to study the SN during spontaneous migraine attacks. Moreover, to isolate the emotional and cognitive aspects of pain processing from the sensory and physiological changes associated with the attack, we aim to also study the SN during an imagery task designed to elicit the experience of a migraine attack in the absence of pain, i.e., during the interictal phase. We hypothesize that such an imagery task should allow us to probe the network’s engagement in pain-related cognitive and emotional processes, while disentangling the nociceptive processing from the other components that take part in the experience of a migraine attack. Several studies have shown that tasks involving the imagination of pain or its suggestion through hypnosis lead to functional changes in the insula and ACC, as well as other regions associated with pain processing, in healthy adults (Derbyshire et al., 2004; Gu & Han, 2007; Ogino et al., 2006; Raij et al., 2005, 2009; Richter et al., 2010). In fact, <u>Eck et al. (2011)</u> showed that during the processing of pain-related words, migraine patients exhibited increased activation of the insula. Furthermore, our previous research indicates that pain imagery can significantly deactivate the ACC, and other pain-processing regions, in migraine patients during the interictal phase, relative to healthy controls (Perdigão et al., 2024).

Therefore, in this study, we examined the dFC of the SN in a group of migraine patients without aura during the ictal phase at rest, as well as during the interictal phase, both at rest and while performing a pain imagery task, using a longitudinal design. For comparison, we also studied a group of healthy controls in three matching conditions, including rest as well as an imagery task of a (non-migraine) head pain experience. This approach allows us to assess the temporal variability of SN connectivity during intrinsic, resting conditions as well as during active, pain-related cognitive engagement. We hypothesize that the pain imagery task may amplify alterations in the temporal dynamics of the SN in migraine patients differently than in controls, even during the interictal phase. Moreover, these patterns may differ from those observed in the ictal phase, as additional components of the pain experience are also involved. By exploring these dynamic connectivity patterns, we aim to uncover the temporal and functional instability of the SN in migraine without aura, and understand its role in the aberrant sensory processing and pain perception, in both spontaneous and stimulus-driven contexts.

## 2. Materials and Methods

This study is part of a broader research project on brain imaging in migraine (MIG_N2Treat). It followed a prospective, longitudinal, within-subject design, and consisted of a comprehensive neuroimaging protocol encompassing arterial spin labelling MRI, diffusion MRI and task-fMRI as well as resting-state fMRI. In this report we focus specifically on the pain imagery task and resting-state fMRI data, while the methods and results of other parts of the protocol are described elsewhere (Domingos et al., 2023; Esteves et al., 2025; Fouto, Henriques, et al., 2024; Fouto, Nunes, et al., 2024; Guadilla et al., 2025; Matoso et al., 2024; Ruiz-Tagle et al., 2025). The research protocol and statistical analysis were not preregistered. The study was approved by the Hospital da Luz Ethics Committee and all participants provided written informed consent in accordance with the Declaration of Helsinki 7th revision.

### 2.1. Participants and experimental design

Migraine patients were recruited by a neurologist during routine consultations at the headache outpatient clinics of Hospital da Luz. Healthy female controls were recruited through social media advertisements targeting the general population and university campuses, ensuring they matched the patient group in gender, age, contraceptive use, and menstrual phase at the time of scanning. We did not conduct a statistical power analysis prior to starting the study. Notwithstanding, we aimed to have a sample size of 15 migraine patients, aligning with previous MRI cohort studies that detected group differences during spontaneous migraine attacks with sample sizes of N=13 (Coppola et al., 2016, 2018) or N=17 (Amin et al., 2018). The inclusion criteria were: being between 18 and 55 years; having at least 9 years of education; speak Portuguese as their first language; being generally healthy (excluding migraine, for the patients), with no diagnosed condition significantly impacting active and productive life or causing life expectancy to be below 5 years. Additionally, patients were required to suffer from low-frequency episodic migraine without aura, diagnosed according to the criteria of the 3rd edition of the International Classification of Headache Disorders (ICHD-3) (International Headache Society, 2018) with menstrual-related migraine attacks. The exclusion criteria were: history or presence of a neurologic condition (excluding migraine, for the patients); history or presence of a psychiatric disorder and/or severe anxiety and/or depressive symptoms as assessed through questionnaires; history or presence of vascular disease; receiving treatment with psychoactive drugs (including anxiolytics, antidepressants, anti-epileptics, and any migraine prophylactics); being pregnant or trying to get pregnant, breastfeeding, being post-menopausal, or using contraception precluding cyclic menses; contraindications for MRI.

We aimed to assess patients during the ictal (M-ict) and interictal (M-inter) phases. For the interictal session, participants were required to be free from pain for at least 48 hours before, with confirmation of the absence of a migraine attack obtained 72 hours post-scan. Healthy controls were assessed in two phases of the menstrual cycle to match the peri-ictal and interictal phases of the patients, yielding respectively the perimenstrual phase (HC-pm) and the post-ovulation phase (HC-postov).

Resting-state (RS) fMRI data were collected (7min, eyes-open) both for M-inter/HC-postov and M-ic/HC-pm. Participants were instructed to keep their eyes open, looking at a black screen, and to not think about anything in particular. For the pain imagery (PI) task, patients (M-inter) performed attack imagery, recollecting the experience of their most severe migraine attack episode, while controls (HC-postov) had to recall their most severe episode of internal pain (e.g. tooth pain). The task consisted of eight 20-s periods of pain imagery alternated with eight 20-s pain relief imagery (∼5min30s), with eyes-closed following verbal instructions. The experimental protocol is schematically represented in Figure 1.

**Figure 1.**
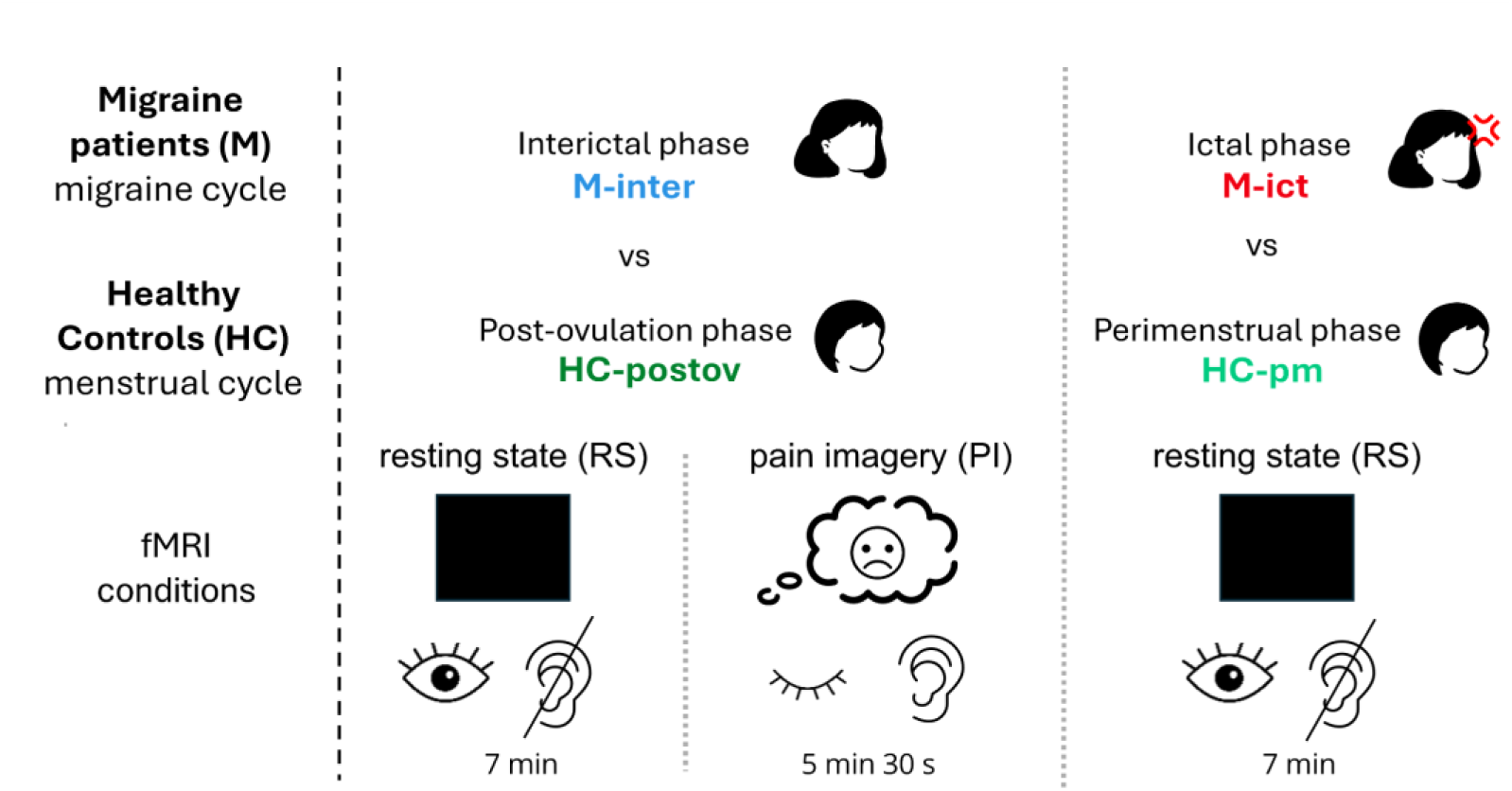
Experimental design: pain-imagery and resting-state fMRI data were acquired for patients and controls in the interictal and post-ovulation phases, respectively (M-inter/HC-postov), whereas only resting-state fMRI data were acquired for patients and controls in the ictal and peri-menstrual phases, respectively (M-ic/HC-pm). Pain imagery (PI): eyes closed, following auditory cues. Resting state (RS): looking at a black screen with eyes open without thinking of anything in particular; no cues.

### 2.2. MRI data acquisition

The MRI data was collected using a 3T Siemens Vida system, with a 64-channel radio frequency (RF) coil. Functional images were acquired with a T2*-weighted gradient-echo Echo Planar Imaging (EPI) with the following parameters: TR/TE=1260/30ms, flip angle = 70°, in-plane generalized autocalibrating partially parallel acquisition (GRAPPA) acceleration factor = 2, simultaneous multi-slice (SMS) factor = 3, 60 slices, 2.2 mm isotropic voxel resolution. T1-weighted structural images were obtained with a magnetization-prepared rapid gradient echo (MPRAGE) sequence, with TR = 2300 ms, TE = 2.98 ms, inversion time (TI) = 900 ms, and 1 mm isotropic voxel resolution. Fieldmap magnitude and phase images were obtained using a double-echo gradient echo sequence (TR = 400.0 ms, TE = 4.92/7.38 ms, voxel size: 3.4 × 3.4 × 3.0 mm^3^, flip angle = 60°). We acquired 262 and 333 fMRI volumes for the pain imagery task and resting state, respectively. Earplugs were provided to reduce acoustic noise exposure, and small cushions were placed around the head to minimize movement. To mitigate acoustic noise exposure and minimize head motion, earplugs were used and small cushions were placed beneath and on the sides of the head of the subject.

### 2.3 MRI data preprocessing

The FMRIB Software Library (FSL)’s tools (Jenkinson et al., 2012) were used to preprocess the MRI data. For the structural images this included nonbrain tissue removal using Brain Extraction Tool (BET) and bias field correction using FMRIB’s Automated Segmentation Tool (FAST). Each structural image was registered to the subject’s functional space (reference volume) as well as the standard MNI space using FMRIB’s Linear Image Registration Tool (FLIRT) and FMRIB’s Non-Linear Image Registration Tool (FNIRT), respectively. fMRI data preprocessing consisted of: EPI distortion correction using FMRIB’s Utility for Geometrically Unwarping EPIs (FUGUE) based on the acquired fieldmap; volume realignment with respect to the middle volume (reference) using (FLIRT), yielding 6 rigid-body motion parameters; and high-pass temporal filtering with a cut-off frequency of 0.01 Hz to remove slow drift fluctuations. Finally, spatial smoothing was performed using SUSAN tool (FSL’s SUSAN), employing a Gaussian kernel with a full-width half maximum of 3.3 mm. The preprocessing pipeline can be found here: https://github.com/martaxavier/fMRI-Preprocessing.

### 2.4 Dynamic functional connectivity (dFC)

For dFC computation and characterization of the temporal dynamics of the SN, we employed the leading eigenvector dynamic analysis (LEiDA) framework (Cabral et al., 2017), previously used to study conditions such as major depressive disorder (Figueroa et al., 2019) and schizophrenia (Farinha et al., 2022). The code for the LEiDA toolbox was publicly available (https://github.com/PSYMARKER/leida-matlab) and was used with MATLAB software version R2023b. The pipeline is illustrated in Figure 2.

**Figure 2.**
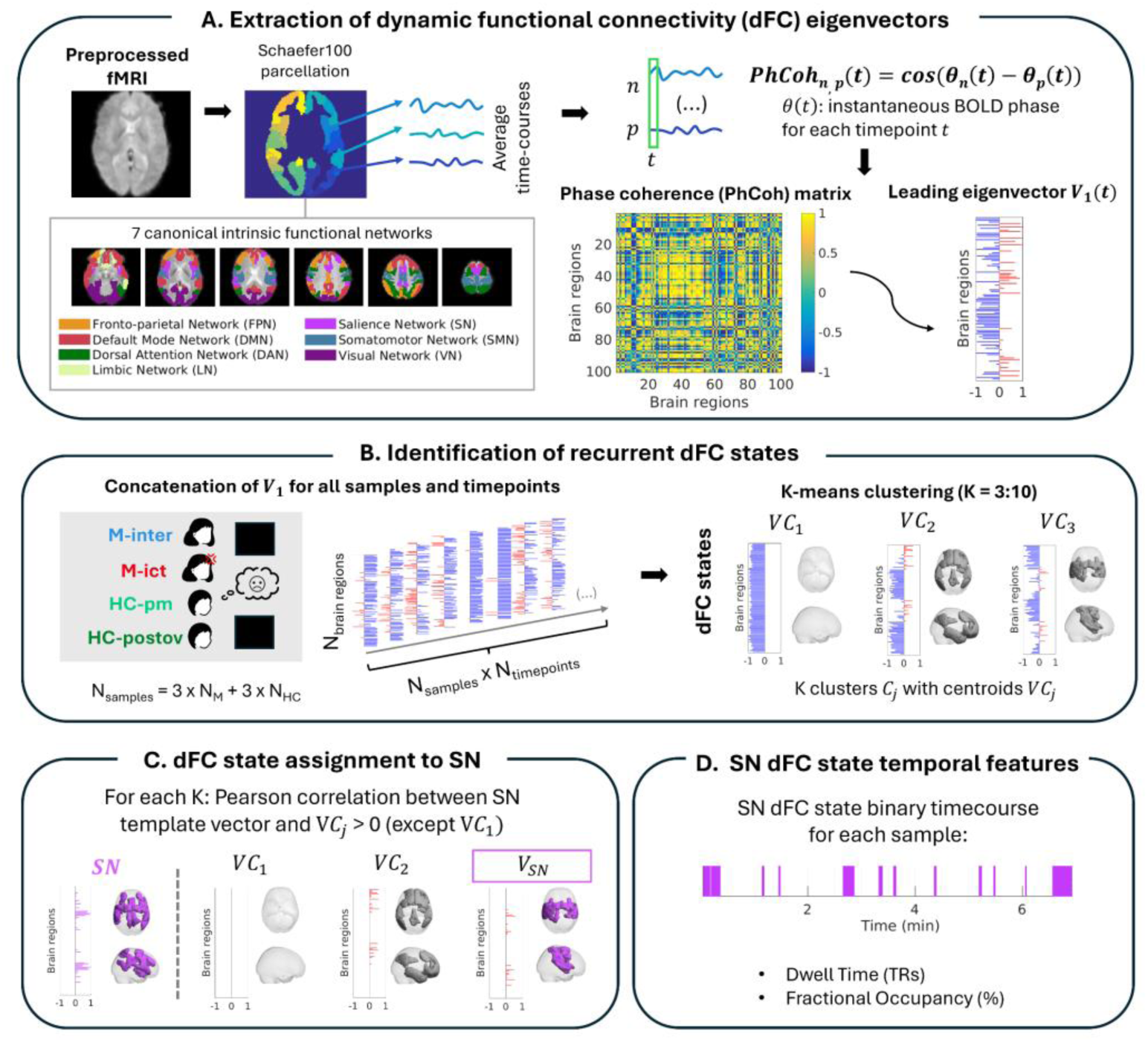
Data Analysis Pipeline: A. Extraction of dynamic Functional Connectivity (dFC) Eigenvectors: Parcellation into cortical regions of intrinsic networks, based on the Schaefer atlas (100 regions, representing 7 networks: salience (SN), fronto-parietal (FPN), default mode (DMN), dorsal attention (DAN), limbic (LN), somatomotor (SMN) and visual (VN). The parcellated data is used for computing the phase coherence (PhCoh) from the instantaneous phase of average time-courses for each pair of regions and the leading eigenvector 𝑉_1_ is extracted from the resulting matrix. **B. Identification of recurrent dFC states:** The 𝑉_1_ for all samples and timepoints are concatenated for a k-means clustering, with the number of clusters ranging from 3 to 10, to identify recurrent dFC states. Each cluster 𝐶_𝑗_ is represented by its centroid 𝑉𝐶_𝑗_ which corresponds to the average of the eigenvectors of the timepoints attributed to that cluster. **C. dFC state assignment to SN:** For each clustering solution with a specific value of K, every cluster centroid 𝑉𝐶_𝑗_is compared to the SN using Pearson correlation and the cluster showing the highest significant correlation is assigned to SN. **D. SN dFC state temporal features:** A binary time course is created for the SN, marking it as dominant only at time points assigned to the cluster identified as the SN. Two metrics describe the SN state dynamics: dwell time (TRs) which is the average duration of a state measure as the number of consecutive fMRI timepoints (the repetition time, TR), and fractional occupancy (%), which represents the probability of the state’s occurrence, computed as the total number of occurrences divided by the total number of timepoints.

#### Extraction of dynamic functional connectivity (dFC) eigenvectors

fMRI data was parcellated by computing region average time courses using Schaefer atlas (Schaefer et al., 2018) with 100 parcels, directly corresponding to the 7 canonical intrinsic functional networks defined by (Thomas Yeo et al., 2011): fronto-parietal network (FPN), default mode network (DMN), dorsal attention network (DAN), limbic network (LN), ventral attention network (VAN), somatomotor network (SMN) and visual network (VN). The time series of each region were demeaned and bandpass-filtered from 0.01 to 0.1 Hz using a 5^th^ order Butterworth filter.

We estimated the individual instantaneous phase of each region (𝜃) for each timepoint by applying the Hilbert Transform to each region time-course. The phase coherence (𝑃ℎ𝐶𝑜ℎ) between regions 𝑛 and 𝑝 for each timepoint 𝑡 was then estimated as:

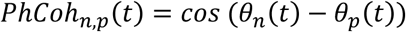

𝑃ℎ𝐶𝑜ℎ values vary between -1 and 1, where -1 indicates that areas 𝑛 and 𝑝 are anti-phase at time 𝑡 and 1 means they are synchronized. For all subjects, we computed it for all pairwise combinations of brain areas (𝑛, 𝑝), with 𝑛, 𝑝 𝜖{1, …,100} at each timepoint 𝑡, with 𝑡 𝜖{2, …,261} for PI or 𝑡 𝜖{2, …,332}. The first and last timepoints for each scan were excluded since they could suffer distortions induced by the Hilbert transform.

Following the LEiDA approach, which concentrates on the principal patterns of 𝑃ℎ𝐶𝑜ℎ matrices by analysing their leading eigenvector, we computed the leading eigenvector for each time point, 𝑉_1_(𝑡), a 100 × 1 vector (reflecting the parcellation into 100 regions) that indicates the main direction of phase alignment across all brain regions, dividing brain regions into two communities based on the sign of its elements. If all elements of 𝑉_1_(𝑡) share the same sign, it reflects a global phase coherence, meaning all regions are grouped into one community. However, if 𝑉_1_(𝑡) contains both positive and negative elements, brain regions are divided into two communities according to their phase coherence. The absolute value of each element of 𝑉_1_(𝑡) indicates how strongly each brain region contributes to its assigned community. We set up 𝑉_1_(𝑡) so that most of its elements are negative, noting that both 𝑉_1_(𝑡) and −𝑉_1_(𝑡) are valid leading eigenvectors. This approach allows us to identify a distinct functional network comprising brain regions with positive elements, representing regions that deviate from the global coherence mode. By employing this methodology, we achieve significant dimensionality reduction while preserving over 50% of the variance in phase coherence across all samples and time points.

#### Identification of recurrent dFC states

After calculating 𝑉_1_(𝑡) from the 𝑃ℎ𝐶𝑜ℎ matrix for each time point, we identified recurring dFC states observed over time. To do this, we combined the leading eigenvectors from all samples, including data from both groups and all three conditions, into a single dataset. We then used k-means clustering to group the leading eigenvectors into clusters, exploring a range of 3 to 10 clusters (𝐾 = 3: 10), representing 8 possible numbers of dFC states. Clustering was conducted by minimizing the cosine distance between eigenvectors within each cluster. The algorithm was run 1,000 times, with a maximum of 500 iterations per run, to minimize the chance of settling on a local rather than a global solution. For each value of 𝐾, the process yielded 𝐾 clusters. Each cluster 𝐶_𝑗_ (𝑗 ∈ {1, …, 𝐾}) was represented by a centroid (𝑉𝐶_𝑗_), a vector of dimension 100 × 1, corresponding to a recurrent dFC state. The centroid was calculated as the average of the leading eigenvectors assigned to that cluster.

#### dFC state assignment to SN

After determining the 𝐾 possible clustering solutions, we identified the centroid that most closely resembled the SN for each number of clusters, i.e., the one with the largest spatial overlap. To achieve this, we created an 𝑁 × 1 vector, 𝑉_𝑆𝑁_, representing the SN, where each element was calculated as the proportion of voxels in each region of Schaefer’s parcellation (with direct matching to Yeo’s intrinsic functional networks) corresponding to the SN, relative to the total number of voxels in that region. We then computed the Pearson correlation between 𝑉_𝑆𝑁_ and each cluster centroid vector, 𝑉𝐶_𝑗_. For this computation, all the negative values of 𝑉𝐶_𝑗_ were set to zero, to compare only the regions belonging to the functional network deviating from the global mode. The centroid with the highest significant correlation with 𝑉_𝑆𝑁_ was designated as representing the SN.

#### SN dFC state temporal features

For each clustering solution, we derived a binary time course for the salience network (SN) state for each subject. This binary time course indicated whether the SN was in the background (when the assigned centroid did not correspond to the SN) or dominant (for time points assigned to the SN centroid). To characterize the SN dynamics, we calculated two parameters: fractional occupancy and dwell time. Fractional occupancy represents the probability of the SN state occurring, defined as the percentage of time points assigned to the SN state. Dwell time refers to the average duration of the SN state, calculated as the mean number of consecutive time points assigned to the SN state across all occurrences. This calculation excludes instances where the state occurs at the very beginning or end of the scan, since it may have already started or may have not finished yet and the duration would not be correctly estimated.

Statistically significant differences between groups during pain imagery and resting state for both sessions were evaluated using a permutation-based t-test. This non-parametric test evaluates group differences by permuting group labels to generate the null distribution, rather than relying on standard parametric distributions. The null distribution is computed separately for each population, and a t-test is performed for each of the 5,000 permutations to compare the groups. For each parameter, we then applied multiple comparisons FDR correction with the Benjamini-Hochberg method (Benjamini & Hochberg, 1995; Martínez-Cagigal, 2021), to control for the number of clustering solutions (𝐾 = 3, …,10) and group comparisons (M-inter vs HC-postov RS; M-inter vs HC-postov PI; M-ict vs HC-pm RS).

## 3. Results

### 3.1 Participants

An initial pool of 29 migraine patients and 60 healthy controls were approached out of which 18 and 16 were included in the study, respectively. Of the 18 patients, one was excluded due to an incidental finding and only 11 migraine patients completed the 2 sessions and were included in the analysis. From the 16 healthy controls, one subject was excluded due to incidental findings, leading to a final sample of 15 healthy controls. Due to technical issues, data acquisition was truncated for two participants. However, given the nature of the analysis and the small sample size, their data were still included. Specifically, for one participant during M-inter PI, 235 out of 262 volumes were acquired, and for another participant during HC-pm RS, 222 out of 333 volumes were acquired. From the 3 conditions of the final groups of 11 migraine patients (age 36.1±8.3 years) and 15 healthy controls (age 30.9±6.8 years), we obtained a total of 78 samples to define the SN states. The flowchart of the recruitment and participant selection is presented in Figure 3.

**Figure 3.**
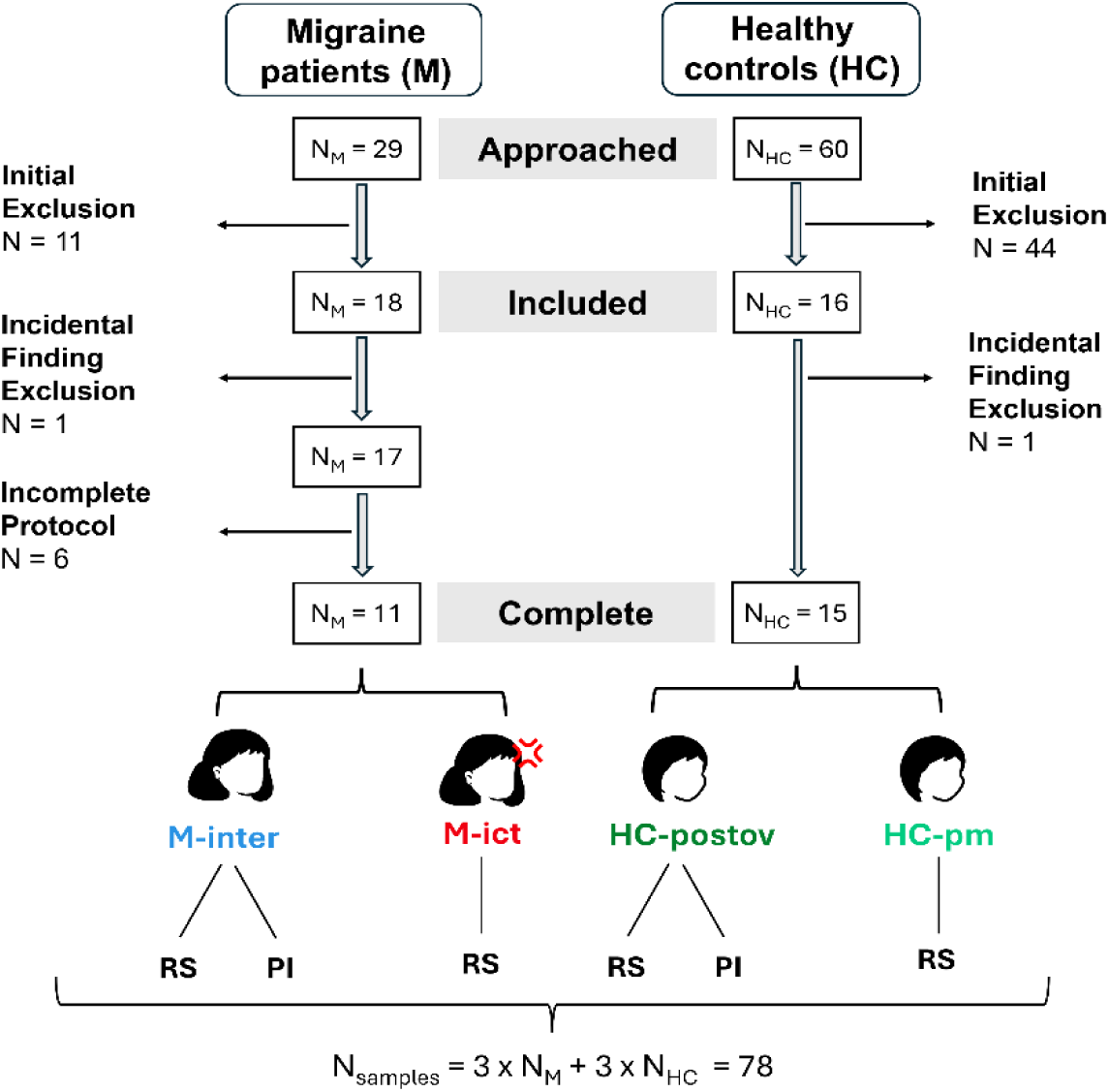
**Flow chart of the study showing the number of participants approached, included, and completing the protocol, with the corresponding fMRI conditions**. Migraine patients in the interictal phase (M-inter) and healthy controls in the postovulation phase (HC-postov) were studied in resting state (RS) and pain imagery (PI) conditions. Migraine patients in the ictal phase (M-ict) and healthy controls in the premenstrual phase (HC-pm) were studied only in RS. With 11 migraine patients and 15 healthy controls each completing three conditions, we collected a total of 78 samples to define the SN states.

### 3.2 dFC analysis

For each clustering solution, we identified the centroids that showed the strongest significant correlation with the SN. Their spatial patterns are presented in Figure 4. From the distribution of the positive elements of the centroid vector, we may observe that most of these elements correspond to regions within the SN. Additionally, the centroid vector highlights the contributions of brain regions from each of the other six canonical intrinsic functional networks defined by Yeo, with other contributions coming mostly from the DAN and SMN. Notably, the clustering solutions were highly consistent across different values of K, reinforcing the robustness of the observed patterns.

**Figure 4.**
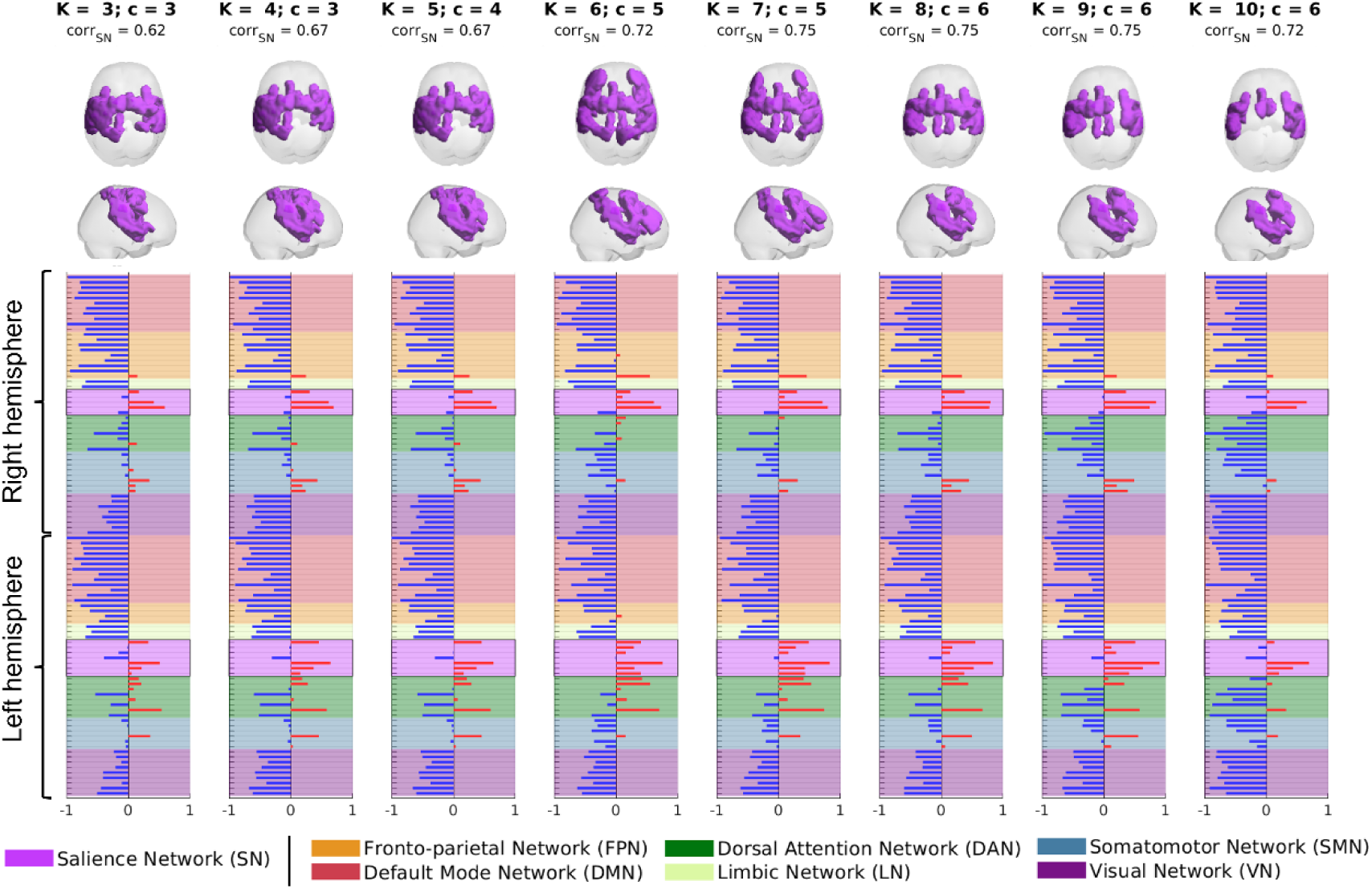
**Centroids that achieved the maximal significant correlation with the SN for each clustering solution**. Top: spatial distribution of positive elements of the centroid vector using axial and sagittal views of the brain volume. Bottom: centroid vector showing the contribution of regions belonging to the seven canonical intrinsic functional networks defined by Yeo to each of the centroids; the regions corresponding to the SN according to Yeo’s definition are highlighted by a box.

For each clustering solution, we make a link with anatomical regions that constitute the SN, by showing the contribution to the centroid attributed to the SN state, in Figure 5. For instance, the ACC and the insula play a key role in the salience network and have been linked to migraine pathophysiology. For 𝐾 = 6 up to 𝐾 = 10, the ACC is represented bilaterally, whereas for other 𝐾’s only the right side shows a contribution. Regarding the insula, it presents a bilateral contribution for all 𝐾’s, being the strongest. In general, the represented regions and the values of their contributions are consistent across 𝐾’s.

**Figure 5.**
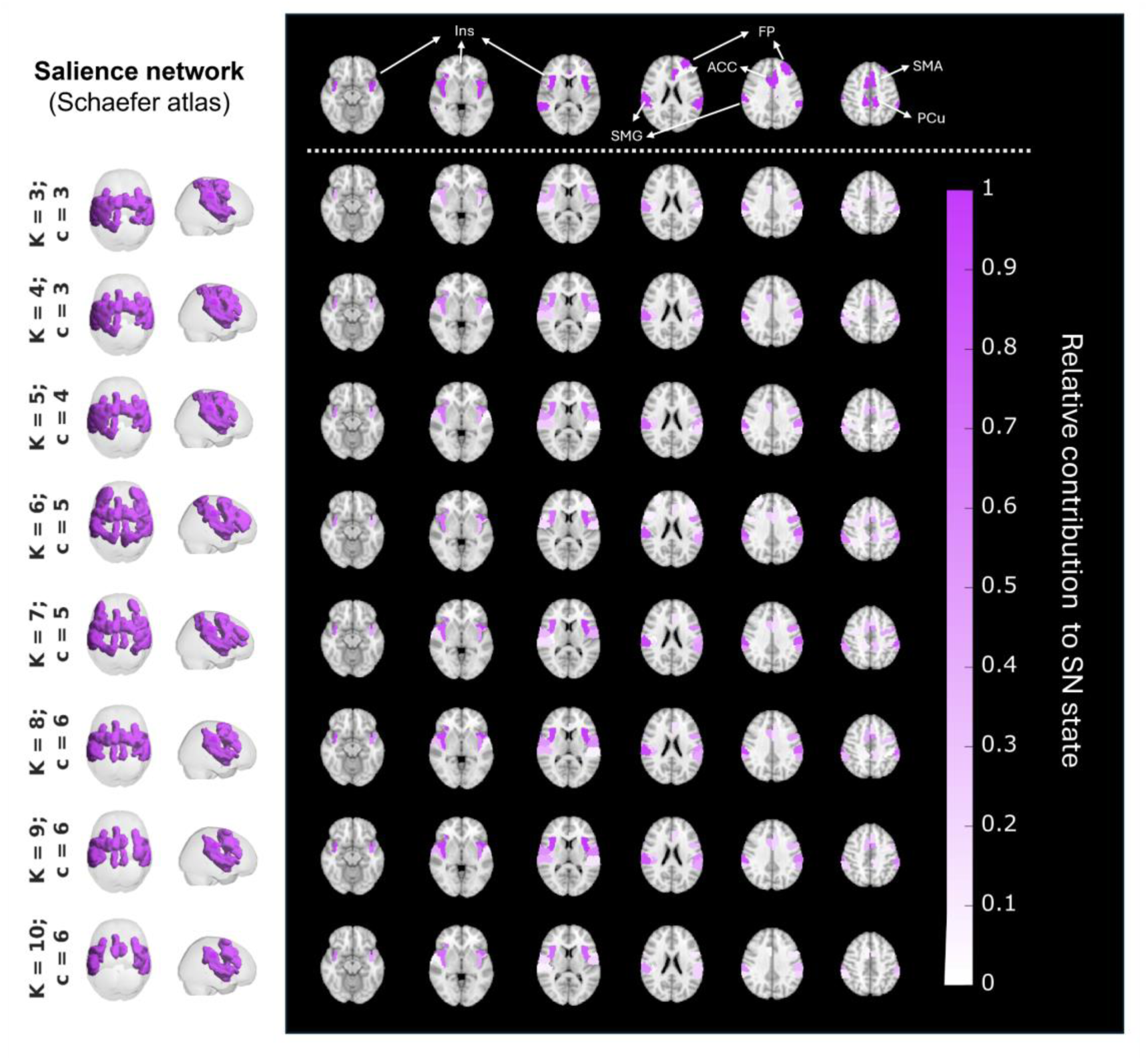
Spatial mapping of the dFC state assigned to the SN. Left) A 3D representation of the vector centroid attributed to the SN. Right) Lightbox view representation of the SN using Schaefer atlas parcels and the centroid vectors that were attributed to SN based on correlation. The first row shows the SN using Schaefer atlas parcels highlighting the parcels that overlap with key SN regions such as the anterior cingulate cortex (ACC) and insula (Ins), along with other regions identified as supramarginal gyrus (SMG), frontal pole (FP), supplementary motor area (SMA) and precuneus (PCu); The other rows show the spatial distribution of centroid vector regions and the corresponding contributions to SN state (only the positive values are represented, since these are the ones associated to the functional network that deviates from the global mode state).

The fractional occupancy and dwell time for all sessions and tasks across 𝐾’s are represented in Figure 6. For the fractional occupancy, there were no significant differences between groups for any of the sessions/tasks. During PI, M-inter showed a significantly lower SN dwell time compared to HC-postov, for 𝐾 = 7, 8, 9. The other clustering solutions present a similar trend, as expected since they correspond to analogous configurations, but did not survive the significance threshold. There were no significant differences for any clustering solution for RS, either for M-inter/HC-postov or M-ic/HC-pm.

**Figure 6.**
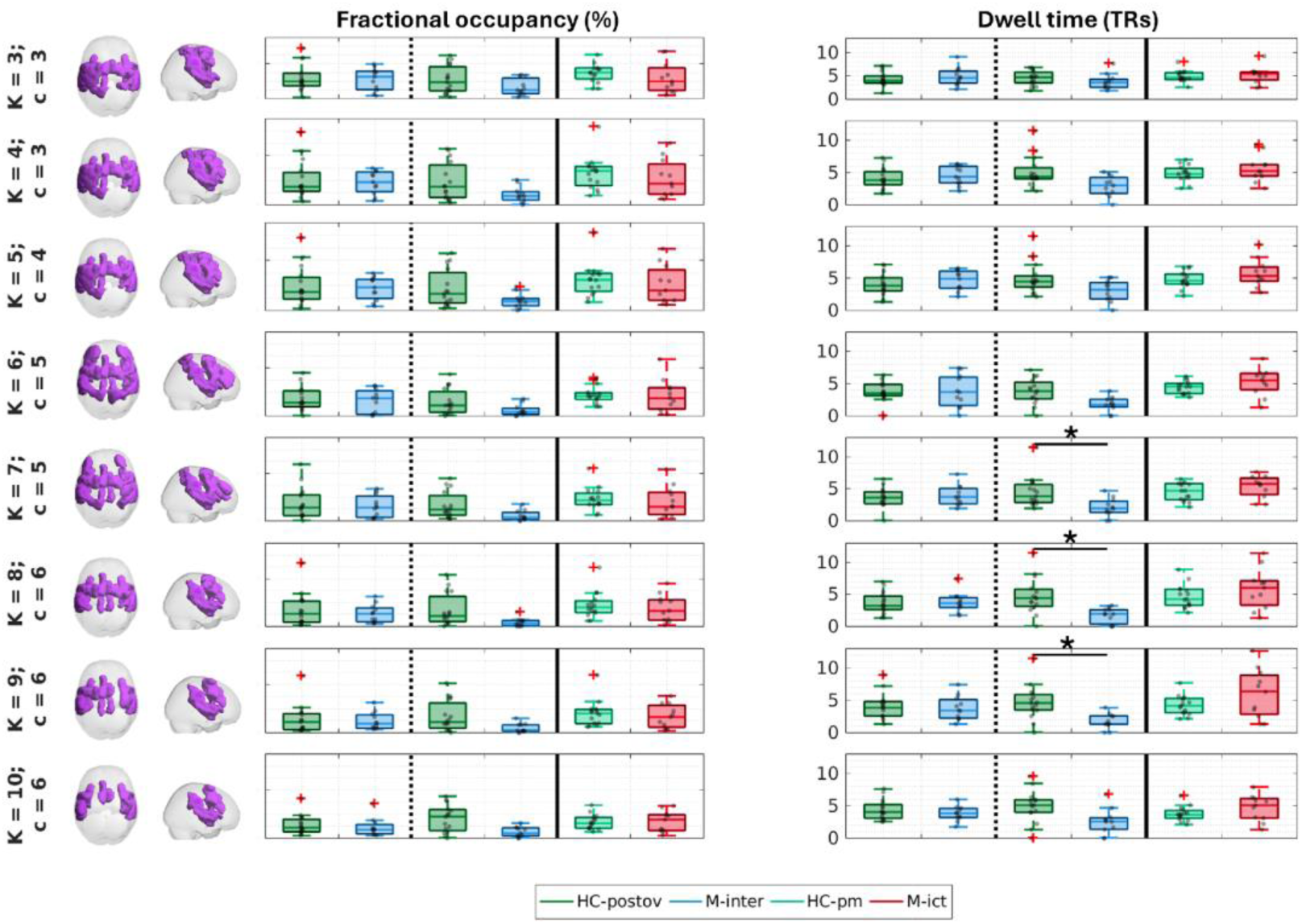
Fractional occupancy (%) and dwell time (TRs) for all sessions and tasks across. 𝑲**’s.** A 3D representation of the vector centroid associated with the SN is displayed on the left to facilitate interpretation. Vertical dashed lines separate tasks acquired within the same session, while vertical solid lines distinguish different sessions. Significant differences found by permutation tests followed by FDR correction are represented by * (corrected p-value < 0.05).

Figure 7 depicts the time-course of the SN for all participants, sessions and tasks, for 𝐾 = 7, the first clustering solution for which there are significant differences regarding the dwell time. Consistently with the shorter dwell time, it can be observed that the SN events tend to be shorter for M-inter PI compared with HC-postov PI, as indicated by the statistical analysis.

**Figure 7.**
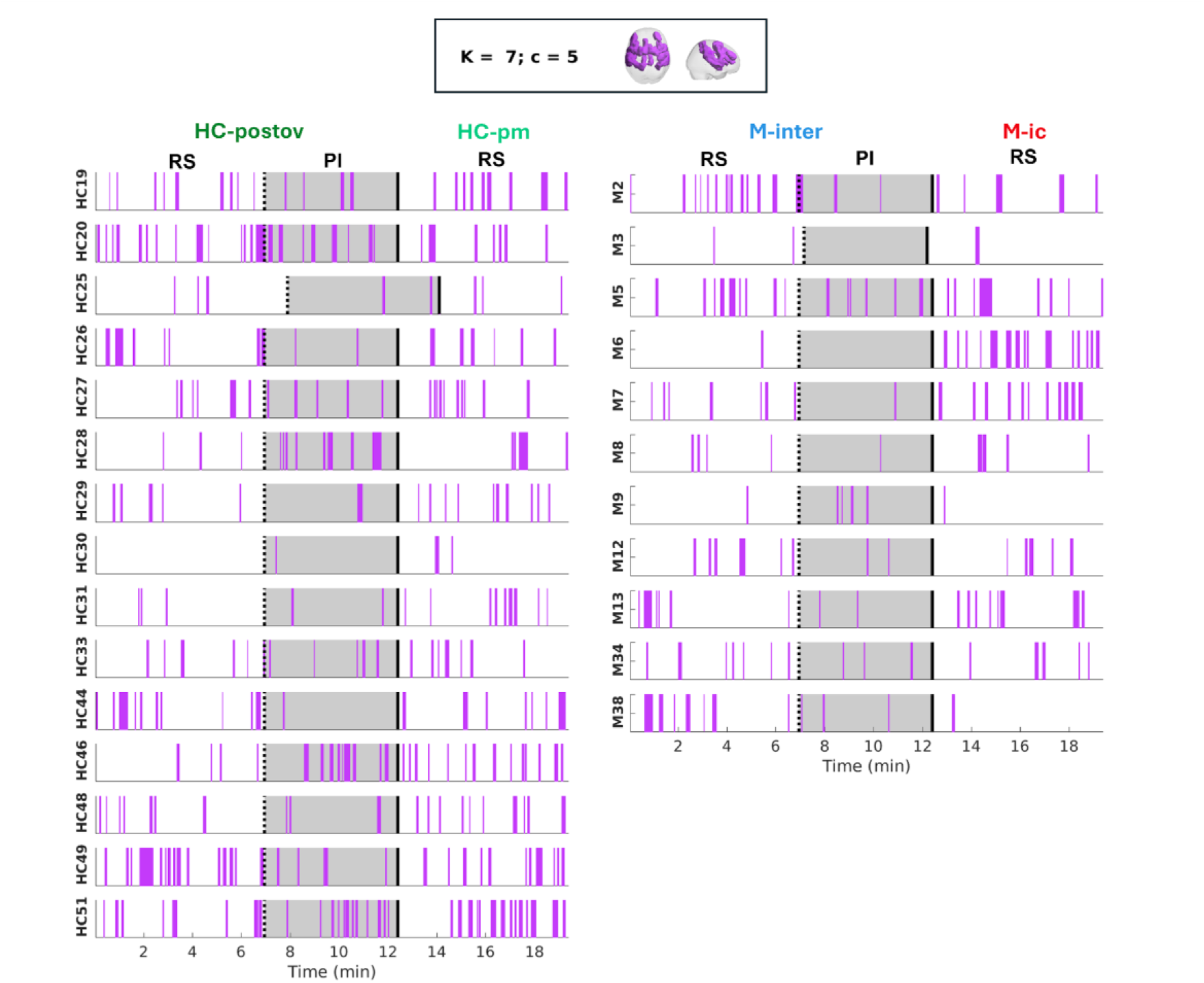
Time-course of SN for all M and HC participants across all sessions and tasks, with K=7. Vertical dashed lines separate tasks acquired within the same session, while vertical solid lines distinguish different sessions. Purple boxes indicate periods of SN activity, with their width reflecting the duration of these periods. Shaded areas represent the PI task, as it was the only task showing significant differences between M and HC. For HC25 and M3, the original fMRI signals were truncated for the HC-pm RS (222/333 volumes) and M-inter PI (235/262 volumes), respectively, resulting in a different time scale.

## 4 Discussion

We found that migraine patients exhibit altered SN dynamics compared to healthy controls, when performing imagery of a previous headache attack during the interictal phase. No differences were observed during resting state in this phase, or even while experiencing a spontaneous migraine attack (in the ictal phase). Dynamic functional connectivity (dFC) analysis revealed that the SN in patients was dominant for shorter periods than in controls, indicating temporal instability. These findings support our hypothesis that migraine is associated with disrupted SN dynamics, with these alterations becoming more pronounced during engagement in an attack imagery task, likely due to the isolation of the emotional and cognitive components of the pain experience from its sensory components.

### Salience network disruption in migraine

Our finding of shorter mean duration of SN occurrences indicates increased temporal instability, which is aligned with the work by <u>Veréb et al. (2020</u>) reporting greater SN temporal instability in migraine patients with aura during the interictal phase. In contrast to <u>Veréb et al. (2020)</u>, we found SN instability in patients without aura and while they were engaged in the imagery of an attack. It is possible that, during the resting state, only migraine patients with aura—who likely exhibit higher cortical excitability—show altered sensory processing that leads to changes in SN dynamics, whereas our use of a pain imagery task enabled us to detect these effects in patients without aura. Our results are also in line with a resting-state electroencephalography (EEG) study reporting that a microstate linked to the SN exhibited a shorter mean duration for patients during the interictal phase compared to healthy controls (Y. Li et al., 2022). Indeed, EEG microstates are thought to reflect the spatiotemporal dynamics of brain networks showing quasi-stable spatial configurations, akin to the fMRI dFC states in our analysis. In contrast with the two previous studies, we did not detect SN impairment during rest but only upon the execution of a pain imagery task. It is possible that impairments were detectable only in this condition due to the heightened demand for inhibitory control, to which this network is linked (Ghahremani et al., 2015).

### Pain imagery and the salience network

This is the first study to employ a pain imagery fMRI paradigm to assess SN connectivity in migraine patients. We have previously investigated brain activation using our proposed pain imagery fMRI paradigm by employing a whole-brain voxel-wise approach (Perdigão et al., 2024). We found that, compared to healthy controls, migraine patients exhibited significantly greater deactivation of the ACC, among other pain-processing regions, indicating a dysregulation of pain inhibition. The ACC is involved in the emotional aspect of pain processing, becoming active when we observe others in pain and when we experience our own pain-related distress. It also helps regulate perceived pain intensity, contributing to the ability to adapt to discomfort and manage suffering in pain responses (Fuchs et al., 2014; Xiao & Zhang, 2018). Given this, it is understandable that even without a painful stimulus this region exhibits dysfunction. The current study further supports it through its involvement in the abnormal dynamics of the SN.

A few fMRI studies have previously analysed brain activation during pain imagery or similar protocols (Derbyshire et al., 2004; Gu & Han, 2007; Ogino et al., 2006; Raij et al., 2005, 2009), reporting the involvement of areas belonging to the SN, such as the ACC, insula, cerebellum, dorsolateral prefrontal cortex and secondary somatosensory cortex. To our knowledge, only <u>Eck et al. (2011)</u> employed a pain imagery paradigm in migraine patients, whereby participants had to create mental images (compared to counting the number of vowels) based on pain-related and non-pain related adjectives. During imagination of pain-related words, patients showed stronger activation in affective pain-related brain regions compared to controls. Interestingly, although the words used in the paradigm were not migraine-specific, five migraine patients and one control referred having imagined a headache as an example of a pain-related word. Despite some similarity with our imagery conditions, the SN connectivity and temporal dynamics were not analysed and therefore the results cannot be directly compared with our current study.

### Areas of the salience network

The insula and ACC are two crucial regions within the SN that contributed to the identification of the SN dFC state in our study. Since pain is a multidimensional experience (Lee et al., 2020), the pain imagery task is expected to primarily trigger the unpleasant emotional component rather than the sensory-discriminative aspect, which aligns with the known roles of the ACC and insula. In fact, observing a noxious stimulus being administered to someone else, or viewing a cue that signals its impending delivery, does not directly activate nociceptors, but it carries a high degree of salience <u>(Iannetti & Mouraux, 2010)</u>.

The insula has been linked to autonomic and somatosensory alterations in migraine, as it is involved in autonomic regulation, interoception and emotion (Menon & Uddin, 2010). The anterior insula is connected to the middle and inferior temporal gyri, the anterior cingulate cortex, and various limbic regions, all of which contribute to its role in emotional regulation. In contrast, the posterior insula is linked to premotor, sensorimotor, supplementary motor areas, and the middle-posterior cingulate cortex, supporting its involvement in sensorimotor integration. The insula has been associated not only with nociception but also with the affective dimension of pain, particularly the unpleasantness of a painful experience (Lee et al., 2020). It is also involved in empathizing with others’ pain (Jackson et al., 2006) and remains active during noxious stimulation even in individuals with congenital insensitivity to pain (Salomons et al., 2016). This aligns with our findings that alterations occur even without painful stimuli; simply imagining a painful and unpleasant experience can lead to observable differences in migraine patients.

### Attack imagery vs. attack experience

Since we could not observe SN disruption during the spontaneous migraine attacks (ictal phase), our results suggest that additional or alternative mechanisms may be active during the experience of an attack compared to the imagery of an attack. These mechanisms might either obscure the contribution of the SN or fundamentally alter its behaviour, by affecting oxygenation, metabolism, and neuronal excitability. For instance, during the ictal phase, heightened cortical excitability or other network-level adaptations could dominate sensory processing, effectively masking the dynamics of the SN observed during the interictal phase or under a pain imagery task. This shift in network dynamics indicates that the migraine attack may involve complex, overlapping processes that differ from those present during periods without an attack.

Another possible explanation for not finding SN disruption in the ictal phase is that sensory processing abnormalities fluctuate throughout the migraine cycle, intensifying in the preictal phase but generally diminishing during the attack (de Tommaso et al., 2014). This could suggest that an adaptive or protective mechanism is active during the interictal phase, leading to an amplification of certain sensory processing components during this period.

Although the proposed attack imagery task clearly does not exactly mimic the experience of a migraine attack, it successfully elicited SN dynamics that was distinct from healthy controls undergoing a matching pain imagery task. These results indicate that this may be a valuable approach for studying functional changes in the brain of migraine patients associated with the recurrent experience of headache attacks. When applied interictally, this attack imagery paradigm appears capable of isolating the emotional and cognitive aspects of the pain experience from other sensory and physiological changes that accompany an attack.

## Limitations

Regarding the pain imagery task, all participants were informed about it before entering the MR scanner and were given the opportunity to select their pain experiences and practice beforehand. However, due to its subjective nature, the content of their imagery may have varied significantly, potentially being less consistent among healthy controls, since they were imagining potentially less specific experiences. Our protocol did not include questionnaires on imagery vividness, which could have provided insight into these variations. Additionally, the potential influence of having eyes open during resting state and closed during the pain imagery task cannot be entirely ruled out, and further data is needed to clarify these effects.

One obvious limitation of our study is the relatively small sample size, which limits statistical power and may not fully capture the diversity of migraine attack experiences and FC patterns. The sample size was constrained by two main factors: our choice of a very specific cohort of migraine patients (menstrual-related low-frequency episodic migraine without aura), and the longitudinal design requiring each patient to be studied over multiple sessions (ictal and interictal). Although the choice of a specific cohort reduces variability between patients eliminating potential confounds, it not only limits the sample size but also restricts the generalizability of our findings to other migraine subtypes. Moreover, despite the strength of the longitudinal design, it should be noted that the data were not always collected within the same migraine cycle for each participant. This may have introduced additional variability due to differences between attacks in the same patient, including pain intensity during the ictal phase.

Another limitation is the use of hormonal contraception by some participants, introducing a potential confound, since hormones influence FC (Dubol et al., 2021) and migraine pathophysiology (Ahmad & Rosendale, 2022; Krause et al., 2021). Unfortunately, restricting the use of hormonal contraception would have further reduced the sample size. Additionally, we did not measure hormonal levels, which renders uncertainty in the definition of the menstrual cycle in women with natural menstrual cycles. Furthermore, the lack of inclusion/exclusion criteria for menstrual cycle regularity or hormone-related disorders, such as premenstrual syndrome or premenstrual dysphoric disorder, further complicates distinguishing migraine-related changes from hormonal influences.

## Conclusions

We employed a case-control longitudinal design to study a group of patients with menstrual-related episodic migraine without aura during the ictal and interictal phases, together with a group of healthy controls in matching phases of the menstrual cycle. We acquired fMRI data during resting state and the execution of a pain imagery task (in the interictal phase) and analysed the brain’s time-varying functional connectivity to identify the dynamics of the SN. We found that, compared to healthy controls, the SN had a significantly shorter lifetime in patients during the pain imagery task, but not during a migraine attack or interictal resting state. Our results support the disruption of the SN in migraine and indicate that pain imagery may be a useful paradigm for isolating the emotional and cognitive aspects of pain and investigating SN dynamics.

